# The alarm pheromone and alarm response of the clonal raider ant

**DOI:** 10.1101/2022.12.04.518909

**Authors:** Lindsey E. Lopes, Erik T. Frank, Zsolt Kárpáti, Thomas Schmitt, Daniel J. C. Kronauer

## Abstract

Ants communicate via an arsenal of different pheromones produced in a variety of exocrine glands. For example, ants release alarm pheromones in response to danger to alert their nestmates and to trigger behavioral alarm responses. Here we characterize the alarm pheromone and the alarm response of the clonal raider ant *Ooceraea biroi*, a species that is amenable to laboratory studies but for which no pheromones have been identified. During an alarm response, ants quickly become unsettled, leave their nest pile, and are sometimes initially attracted to the source of alarm, but ultimately move away from it. We find that the alarm pheromone is released from the head of the ant and identify the putative alarm pheromone as a blend of two compounds found in the head, 4-methyl-3-heptanone and 4-methyl-3-heptanol. These compounds are sufficient to induce alarm behavior alone and in combination. They elicit similar, though slightly different behavioral features of the alarm response, with 4-methyl-3-heptanone being immediately repulsive and 4-methyl-3-heptanol being initially attractive before causing ants to move away. The behavioral response to these compounds in combination is dose-dependent, with ants becoming unsettled and attracted to the source of alarm pheromone at low concentrations and repulsed at high concentrations. While 4-methyl-3-heptanone and 4-methyl-3-heptanol are known alarm pheromones in other more distantly related ant species, this is the first report of the chemical identity of a pheromone in *O. biroi*, and the first alarm pheromone identified in the genus *Ooceraea*. Identification of a pheromone that triggers a robust, consistent, and conserved behavior, like the alarm pheromone, provides an avenue to dissect the behavioral and neuronal mechanisms underpinning chemical communication.

## INTRODUCTION

Many animals use alarm signals to alert conspecifics to danger, yet the modality of these signals varies. Some animals use auditory alarm calls, while others use visual or chemosensory signals or a combination of signals from multiple modalities (Hollén and Radford 2009). Ants use alarm pheromones to alert their nestmates to danger (Blum 1969; Hölldobler and Wilson 1990; Vander Meer and Alonso 1998).

Alarm pheromones are generally present in detectable quantities in ants, which makes them tractable for chemical characterization (Blum 1969). Chemical components of the alarm pheromone are most commonly synthesized in the mandibular gland, Dufour’s gland, and/or poison gland. Some alarm pheromones are multicomponent, and sometimes a single chemical compound is sufficient to induce a behavioral effect (Blum 1969). Components of the alarm pheromone are generally volatile, low molecular weight compounds, often terpenes, ketones, and aldehydes (Amoore et al. 1969; Blum 1969; Brückner et al. 2018; Pokorny et al. 2020; Han et al. 2022).

Alarm behaviors, which are triggered by alarm pheromones, are robust, innate, and present across ant species. However, alarm behaviors can be difficult to quantify, because “alarm behavior” is often used to describe diverse behavioral responses to danger. Broadly, alarm behaviors are characterized by increased alertness, sometimes in combination with moving away from or towards the dangerous stimulus (Hölldobler and Wilson 1990). Some behaviors that occur as part of alarm responses, such as trail-following, aggression, and recruitment, also occur independently from alarm responses, further complicating the characterization of alarm behavior in ants (Hölldobler and Wilson 1990).

Alarm behaviors are broken down into two broad categories, panic and aggressive alarm responses (Blum 1969). Behaviors associated with panic alarm responses include rapid non-directional movement, moving away from the source of alarm, and sometimes nest evacuations. Behaviors associated with aggressive alarm responses include rapid movement towards the source of alarm, assumption of defensive postures, and attack of foreign objects. Different alarm behaviors can be elicited depending on the species of ant, the blend and concentration of pheromone components, and the context in which alarm pheromone is released (Blum 1969; Hölldobler and Wilson 1990; Vander Meer and Alonso 1998).

Many ants use multicomponent alarm pheromones, which can include chemical components that elicit different behavioral responses (Blum 1969; Hölldobler 1995). In the case of the carpenter ant, *Camponotus obscuripes*, alarm behavior is triggered by a blend of formic acid, which is repulsive and released from the poison gland, and *n*-undecane, which is attractive and found in the Dufour’s gland (Fujiwara-Tsujii et al. 2006). Alarm behavior in these ants has different intensities which may depend, at least in part, upon the ratio and volatility of components in the alarm pheromone. In the weaver ant, *Oecophylla longinoda*, alarm pheromone is released from the mandibular gland, which has over thirty compounds present in detectable quantities (Bradshaw et al. 1975). Major chemical components of the mandibular gland secretion, 1-hexanol and hexanal, trigger alert and attraction, and less volatile minor components are thought to be markers for attack (Bradshaw et al. 1975, 1979).

Situation-specific differences in alarm responses have also been described for some ant species. When *Pogonomyrmex badius* harvester ants are alarmed, they can have high or low intensity alarm responses. The principal alarm pheromone of these ants is a ketone, 4-methyl-3-heptanone, that is released by the mandibular gland (McGurk et al. 1966). The low intensity behavioral response includes an increase in locomotion, antennae and head waving, movement in loops and circles, and periodic lowering of the gaster to the ground. During high intensity behavioral responses, there is more locomotion, tighter circling, mandible opening, and less antennae/head waving. High intensity behavioral responses are more likely to occur close to the nest, while lower intensity responses generally occur farther from the nest in the foraging area (Wilson 1958; McGurk et al. 1966).

Here, we describe the alarm behavior of the clonal raider ant *Ooceraea biroi*, identify its putative alarm pheromone, and validate it using electroantennography and behavioral assays. *O. biroi* is a queenless species where all workers reproduce asexually and clonally (Ravary and Jaisson 2004; Kronauer et al. 2012), making it genetically accessible (Trible et al. 2017). *O. biroi* is thus a promising system to study the genetic and neuronal underpinnings of social behavior. However, so far, no pheromones have been identified in this species.

## METHODS AND MATERIALS

### Colony maintenance

Stock colonies of clonal line B *O. biroi* ants, a lineage originally collected in Jolly Hill, St. Croix (Kronauer et al. 2012), were maintained in the laboratory at 25°C in Tupperware containers with a plaster of Paris floor. *O. biroi* colonies are phasic and alternate between reproductive and brood care phases. As described previously (Trible et al. 2017), during the brood care phase, stock colonies were fed with frozen *Solenopsis invicta* brood. For each round of experiments, 12-15 colonies of mixed age ants were established without brood from a single stock colony while the ants were in brood care phase.

### Experimental arena and colony setup

Behavior experiments were conducted in arenas constructed from cast acrylic with a plaster of Paris floor. Each arena was made of four layers; the base layer, a layer of plaster of Paris, a layer with two cut out areas separated by a tunnel, and a top layer of clear acrylic with lids. The arenas were 7 cm x 2 cm in total, with a 2 cm x 0.3 cm tunnel separating two 2.5 cm x 2 cm areas (**Fig S1**). Each area contained a 0.5 cm x 2 cm “stimulus chamber” separated from a 2 cm x 2 cm “nest chamber” by a cast acrylic mesh wall. The wall was laser cut from 0.8 mm thick cast acrylic with multiple holes with a diameter of ~50 μm, as described previously (Chandra et al. 2021). The clear acrylic lids of the nest and stimulus chambers were separate, allowing the experimenter to open the stimulus chamber without opening the nest chamber, thereby decreasing the likelihood of alarming the ants due to increased airflow. The floor of the tunnel was covered with vapor-permeable Tyvek paper to dissuade ants from forming their nest in the tunnel, as described previously (Chandra et al. 2021).

In each arena, 30 ants were introduced without any brood. For the live ant and crushed body experiments (see below), 30 additional ants from the same stock colony were kept in a separate Petri dish with a plaster of Paris floor. These ants were used as the stimulus during experiments.

Ants were fed every 1-2 days with *S. invicta* brood and allowed to lay eggs. About 7-10 days after introducing ants into the arenas, once ants had settled, laid eggs, and clustered into a tightly packed pile (the “nest”) in one of the two nest chambers (**Fig S1**), we began behavioral experiments.

### Behavioral assays with agitated ants and body parts

On each day of behavioral experiments, the acrylic lid of the stimulus chamber on the same side as the ants’ nest was removed. Alarm arenas were placed into an enclosed container with controlled light-emitted diode lighting (STN-A40K80-A3A-10B5M-12V; Super Bright LEDs Inc, Earth City, Missouri, USA) and videos were taken using webcams (Item #V-U0021; Logitech V-U0021, Lausanne, Switzerland). Images were taken at a rate of 10 frames per second and 2,592 × 1,944-pixel resolution. Prior to adding a stimulus, baseline behavior was recorded for 5 minutes. Behavior was recorded for another 5 minutes after exposure to the stimulus.

For the live alarmed ant experiments, 4 minutes and 30 seconds into the recording, an ant from a separate dish was agitated by repeatedly picking her up and putting her down with forceps. This “alarmed ant” was then added to the open stimulus chamber at 5 minutes into the recording. In control experiments, a folded piece of filter paper was added into the stimulus chamber instead.

For the body part experiments, the head of an ant was removed using forceps. After recording baseline activity for 5 minutes, the head or headless body of the ant was crushed using forceps and then quickly added to the open stimulus chamber.

After each assay, the arena was removed, and the behavioral recording box was left open for 1-5 minutes prior to adding the next arena. The total number of ants in the arena was manually counted. Behavioral assays were performed every other day to allow ants to re-settle into a nest after being alarmed. After each behavioral assay, ants were fed with frozen *S. invicta* brood. The following day, the remaining food was removed.

### Chemical analysis

To identify candidate alarm pheromone components from the head, we placed 5 dissected *O. biroi* heads, mesosomas, gasters, or full bodies in a glass-wool-packed thermodesorption tube and added it in the thermodesorber unit (TDU; TD100-xr, Markes, Offenbach am Main, Germany). The thermodesorption tube was heated up to 260°C for 10 minutes. The desorbed components were transferred to the cold trap (5 °C) to focus the analytes using N_2_ flow in splitless mode. The cold trap was rapidly heated up to 310 °C at a rate of 60 °C per minute, held for 5 minutes, and connected to the gas-chromatography/mass-spectrometry unit (GC-MS, Agilent 7890B GC and 5977 MS, Agilent Technologies, Palo Alto, CA, USA) via a heated transfer line (300 °C). The GC was equipped with a HP-5MS UI capillary column (0.25 mm ID × 30 m; film thickness 0.25 μm, J & W Scientific, Folsom, CA, USA). Helium was the carrier gas using 1.2874 ml/min flow. The initial GC oven temperature was 40 °C for 1 minute, then raised to 300 °C at 5 °C per min, where it was held for 3 minutes. The transfer line temperature between GC and MS was 300 °C. The mass spectrometer was operated in electron impact (EI) ionization mode, scanning m/z from 40 to 650, at 2.4 scans per second. Chemical compounds were first identified using the NIST library and later confirmed with co-elution of synthetic 4-methyl-3-heptanone (Pfaltz and Bauer M19160) and 4-methyl-3-heptanol (Sigma Aldrich M48309). Compounds eluting after 30 minutes were excluded from the analysis due to lack of volatility.

To estimate the absolute amount of alarm pheromone in one ant head we injected 75ng of 4-methyl-3-heptanone and 25ng of 4-methyl-3-heptanol into a glass-wool-packed thermodesorption tube and analyzed it using the same method as was used for the ant body parts. We then calculated the area under the peak for the two compounds and compared it to the area under the peak of the sample of 5 *O. biroi* heads (Fig. 2).

**Figure 1.**
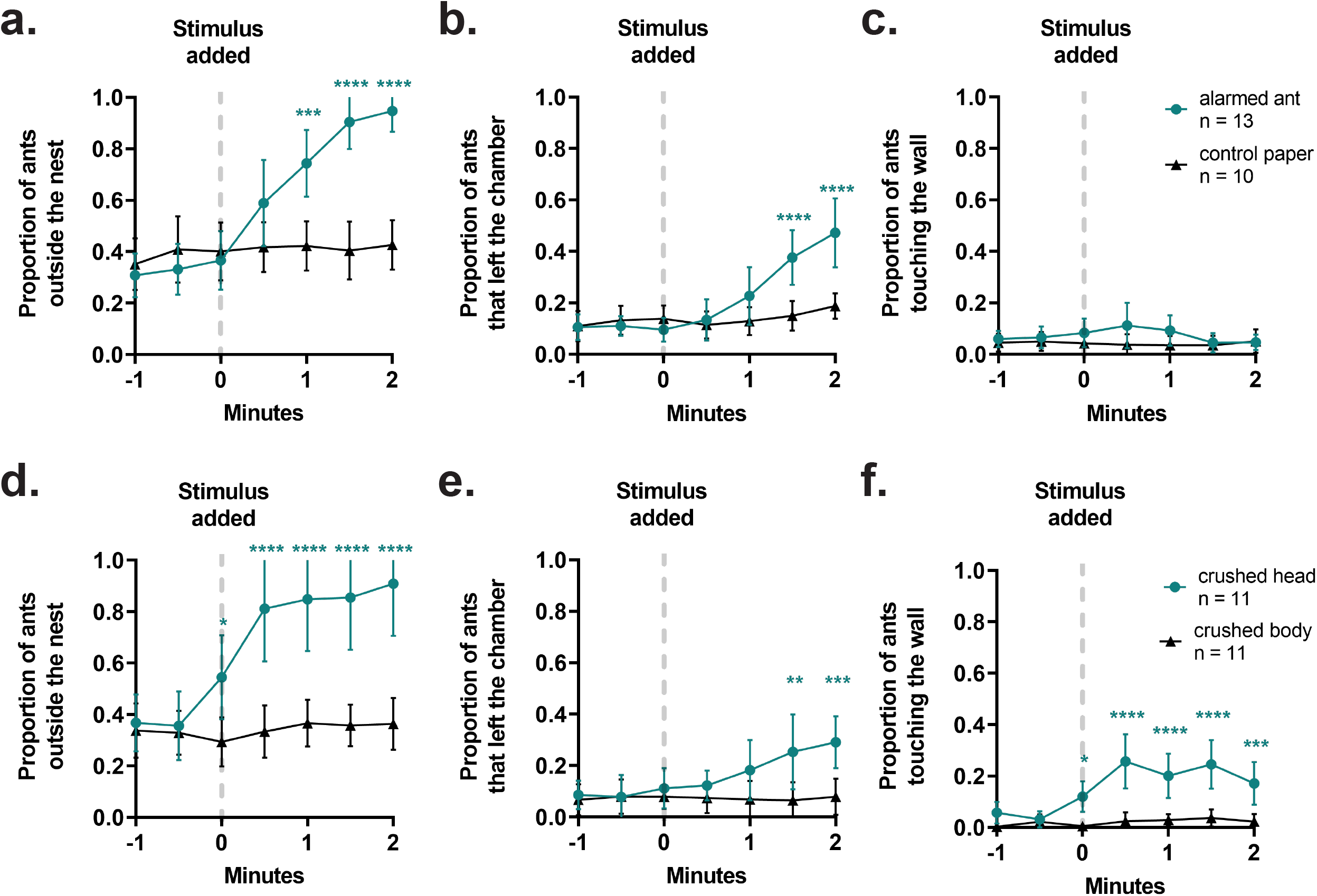
Characterization of alarm behavior and localization of alarm pheromone in *O. biroi*. Quantification of features of the behavioral response of *O. biroi* colonies to a live alarmed ant (a-c) and crushed body parts of an ant (d-f). Data are included from 1 minute prior to adding the stimulus until 2 minutes after. Individual datapoints indicate means and error bars denote 95% confidence intervals. Sample sizes represent replicate colonies tested. Statistical comparisons were performed using a 2-way repeated measures ANOVA with Šidák’s multiple comparisons test to compare individual timepoints. *p<0.05, **p<0.01, ***p<0.001, ****p<0.0001.

**Figure 2.**
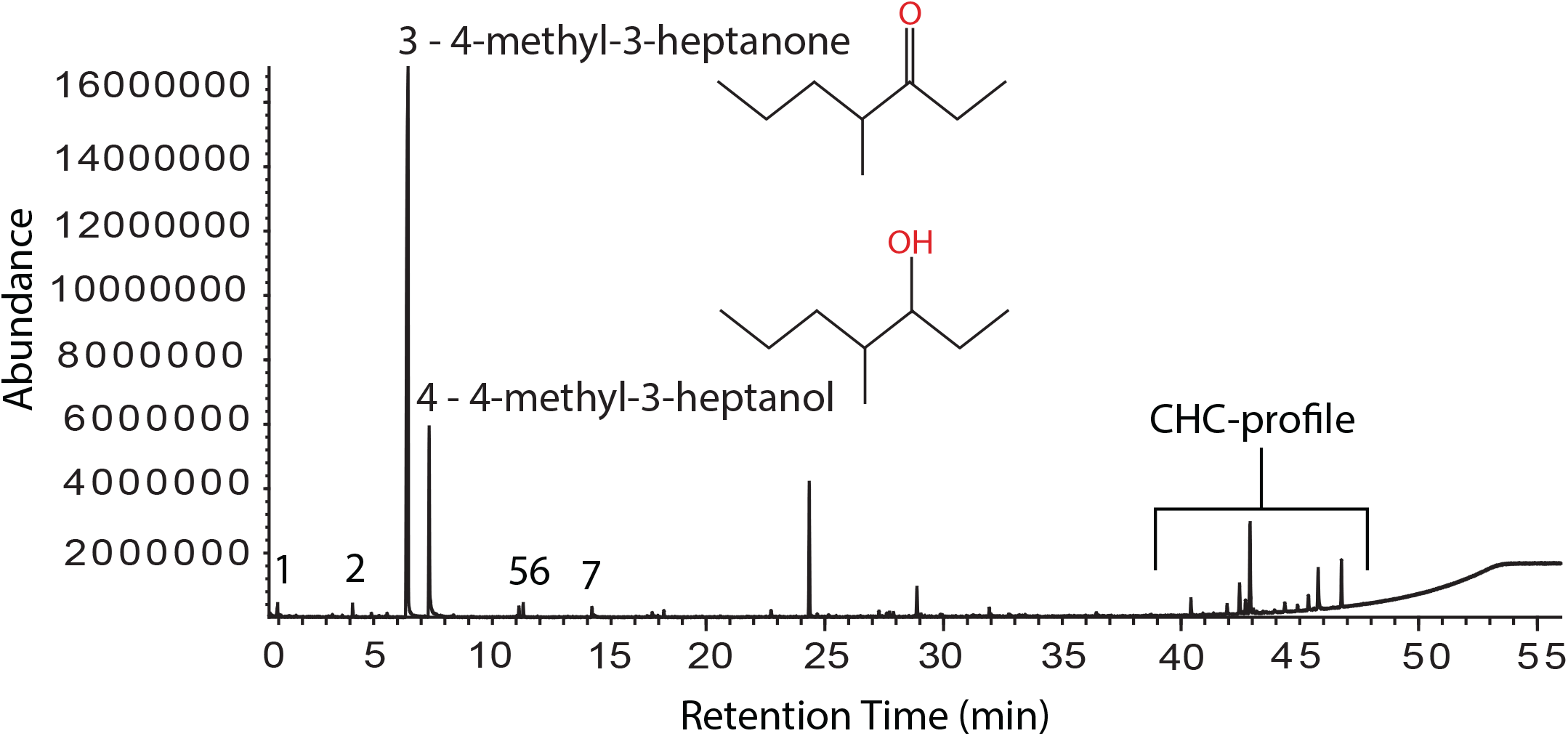
Chemical compounds in the head. Gas-chromatographic representation of one sample of 5 pooled heads. A detailed list of all chemical compounds found in the head (the numbered peaks) is provided in Table 1.

### Chemicals

96% 4-methyl-3-heptanone (mixture of stereoisomers) was purchased from Pfaltz and Bauer (Item #: M19160) and ?99% 4-methyl-3-heptanol (mixture of stereoisomers) was purchased from Sigma-Aldrich (Item #: M48309). Compounds were freshly diluted on each day of behavioral experiments. Dilutions were made using 100% pentane purchased from Sigma-Aldrich (Item #: 236705) as the solvent and diluted compounds were kept in glass vials with a silicone/PFTE magnetic screw cap (Gerstel 093640-079-00) to prevent evaporation.

### Electroantennographic recordings with chemical compounds

For the electroantennographic (EAG) recordings, the head of *O. biroi* was excised and inserted into a glass capillary (ID 1.17 mm, Syntech, Kirchzarten, Germany) filled with Ringer solution (Ephrussi and Beadle, 1936) and attached to the reference silver electrode. The tip of the antenna was inserted into the recording glass electrode, which was also filled with Ringer solution. The antennal signal was 10 times amplified, converted to a digital signal by a high input impedance DC amplifier interface (IDAC-2, Syntech) and recorded with GC-EAD software (GC-EAD 2014, version 1.2.5, Syntech). Synthetic pheromone candidates (4-methyl-3-heptanone and 4-methyl-3-heptanol) were applied at 1, 10 and 100 μg doses on a square filter paper (1×1 cm), which was inserted into a Pasteur-pipette. Stimulus air (2 liters/min) was led into a constant, humidified and charcoal filtered air stream (2 liters/min) using a Stimulus Controller CS-55 (Syntech). Each stimulation was given for 0.5 s. *N*-pentane, the solvent of the components, was used as a control stimulus. Three different doses of the pheromone candidates and the control were randomized and tested on 9 antennae.

Statistical analyses were performed in GraphPad Prism 9.0. To account for differences between individual antennae, we analyzed data using mixed-effects analysis with a Geisser-Greenhouse correction which treats each antenna as a random factor in the model. This method uses a compound symmetry covariance matrix and is fit using Restricted Maximum Likelihood (REML), and results can be interpreted like repeated measures ANOVA in cases with missing values. We then compared each compound/ amount to the pentane solvent, using Dunnett’s multiple comparisons test.

### Behavioral assays with chemical compounds

The alarm behavioral assay was performed as described above. Four minutes and thirty seconds after starting the experiment, 50 μl of the compound diluted in pentane or pentane alone (vehicle control) were added onto a small square of filter paper (~1 cm^2^) using a syringe. The pentane was allowed to evaporate for 30 seconds and then the paper was folded and placed into the open stimulus chamber 5 minutes into the recording.

### Behavioral data analysis

Behavioral recordings were analyzed by hand based on three metrics of interest, (1) number of ants outside the nest pile, (2) number of ants outside the nest chamber, and (3) number of ants touching the mesh wall of the stimulus chamber. A single frame of the recording was scored according to these metrics every 30 seconds for the duration of the 10 minute recording. Ants were scored as being outside the nest pile if they were not touching any other ant within the region of the nest pile (**Fig S1**). Ants were scored as being outside the nest chamber if at least half of their body was outside of the chamber that contained the nest. Ants were scored as touching the mesh wall if any part of their body was in contact with the mesh wall. When nests were touching the wall prior to adding a stimulus, only ants that were outside the nest pile were counted as touching the wall. The proportions of ants outside the nest pile, ants that left the nest chamber, and ants touching the mesh wall were calculated by dividing the number of ants performing each behavior by the total number of ants in the arena. Assays were excluded from further analysis if the average proportion of ants outside the nest pile prior to adding the stimulus was over 0.5, if there was more than one nest pile, or the nest pile was in the tunnel.

Statistical analyses were performed in GraphPad Prism 9.0. We limited the formal statistical analyses to the time window starting 1 minute before the addition of the stimulus and ending 2 minutes after the stimulus had been added, because this was when relevant behavioral changes occurred. These analyses are fully consistent with behavioral dynamics across the entire time course (**Fig S2&3**). To evaluate the effect of the stimulus over time, we performed a two-way repeated measures ANOVA. The effect of the stimulus at each timepoint was then analyzed with Šídák’s multiple comparisons test when comparing two stimuli (live alarmed ant and crushed body part assays) and Dunnett’s multiple comparisons test when comparing to the control stimulus (candidate alarm pheromone assays). To compare features of the behavioral response to 4-methyl-3-heptanone, 4-methyl-3-heptanol, and the blend of compounds, we calculated the area under the curve in the two minutes following addition of each stimulus. Because the different synthetic compounds were tested in different sets of experiments, we also compared the vehicle controls across these three sets of experiments to confirm there were no significant differences in ants outside the nest pile, ants that left the nest chamber, and ants touching the stimulus chamber wall. To evaluate the effect of the compound/blend across concentrations, we performed two-way ANOVAs with Tukey’s multiple comparisons tests on the areas under the curves.

## RESULTS

### Characterization of the clonal raider ant alarm behavior

We began characterizing the alarm behavior of *O. biroi* by quantifying features of the behavioral response of colonies to a live alarmed ant (**Video S1**). Prior to adding the alarmed ant to the stimulus chamber, ants were primarily settled in a nest pile on one side of the arena. When the live alarmed ant was added, ants left the nest pile (**Fig 1a, Table S1**), and some also left the chamber that initially contained the nest pile (**Fig 1b, Table S1**). This response was absent when adding a control, a piece of paper meant to mimic the potential increase in airflow from opening and adding an item into the stimulus chamber. Ants were not attracted to the live alarmed nestmate (**Fig 1c, Table S1**).

### Localization of the clonal raider ant alarm pheromone

To determine from where in the body the alarm pheromone is released, we tested the behavioral response of colonies to crushed ant heads or to crushed bodies without heads (**Video S2**). We hypothesized that the alarm pheromone is coming from the mandibular glands within the head of the ant, or from the Dufour’s and/or poison gland in the abdomen based on studies in other ant species (Blum 1969). In response to crushed heads, *O. biroi* colonies displayed alarm responses like those elicited by live alarmed ants, with an increase in ants outside the nest pile (**Fig 1d, Table S1**) and ants leaving the nest chamber (**Fig 1e, Table S1**). Interestingly, unlike in the response to live alarmed ants, ants were initially attracted to the crushed heads (**Fig 1f, Table S1**). No response was evident when the ants were exposed to crushed bodies without the head (**Fig 1d–f, Table S1**). These data indicate that the volatile compound(s) found in the head of the ant are necessary and sufficient to induce an alarm response, meaning that the alarm pheromone is likely released from the head of *O. biroi*. We therefore conducted GC-MS analyses of the head contents and identified two main compounds, 4-methyl-3-heptanone (80.1% of the head contents) and 4-methyl-3-heptanol (16.3% of the head contents) (**Fig 2, Table 1**). Both compounds only occurred in the head of the ants (**Fig 2, Fig S4**). Based on data from a single sample, we estimate there to be 3.21 μg of 4-methyl-3-heptanone and 0.65 μg of 4-methyl-3-heptanol in the head of an *O. biroi* worker.

**Table 1.** Chemical compounds found in the head contents. Chemical compounds in bold were tested as alarm pheromones. Five heads were pooled per sample run in the GC-MS coupled to a thermodesorption unit as shown in Fig 2.

| Peak # | Compound | Ret.<br>Index | Relative abundance [%] |
| --- | --- | --- | --- |
| 1 | Acetic acid | 702 | 0.94 |
| 2 | 4-Methyl-3-hexanone | 845 | 0.72 |
| 3 | **4-Methyl-3-heptanone** | 940 | 80.08 |
| 4 | **4-Methyl-3-heptanol** | 973 | 16.28 |
| 5 | Undecane | 1105 | 0.68 |
| 6 | Nonanal | 1111 | 0.87 |
| 7 | Decanal | 1213 | 0.43 |

If 4-methyl-3-heptanone and 4-methyl-3-heptanol make up the *O. biroi* alarm pheromone, then ants should be able to detect both compounds with their antennae. To test this, we utilized EAG recordings and found that both compounds were detected (REML mixed effects model difference between treatments p=0.0022; **Fig S5**, **Table S2**).

### Behavioral responses to candidate alarm pheromone components

To determine if 4-methyl-3-heptanone and 4-methyl-3-heptanol are behaviorally active and can trigger an alarm response, ants were exposed to both compounds individually and in combination at two doses, 260 μg and 2600 μg. Pentane was used as the solvent and vehicle control for all assays.

In response to 4-methyl-3-heptanone, ants rapidly left the nest pile (**Fig 3a, Table S3, Video S3**) and left the nest chamber in a dose-dependent manner (**Fig 3b, Table S3)**. There was a small but significant increase in the proportion of ants touching the wall after exposure to both concentrations of 4-methyl-3-heptanone (**Fig 3c, Table S3**). 260 μg 4-methyl-3-heptanone attracted ants for slightly longer than 2600 μg, and by 1.5 minutes after addition of 4-methyl-3-heptanone there was no longer a significant difference between either amount of compound and the vehicle control. However, the increase in the average proportion of ants attracted to 4-methyl-3-heptanone was small. The increase in the average proportion of ants that moved away from the stimulus was much greater, indicating that this compound is mostly repulsive to the ants, especially at high doses.

**Figure 3.**
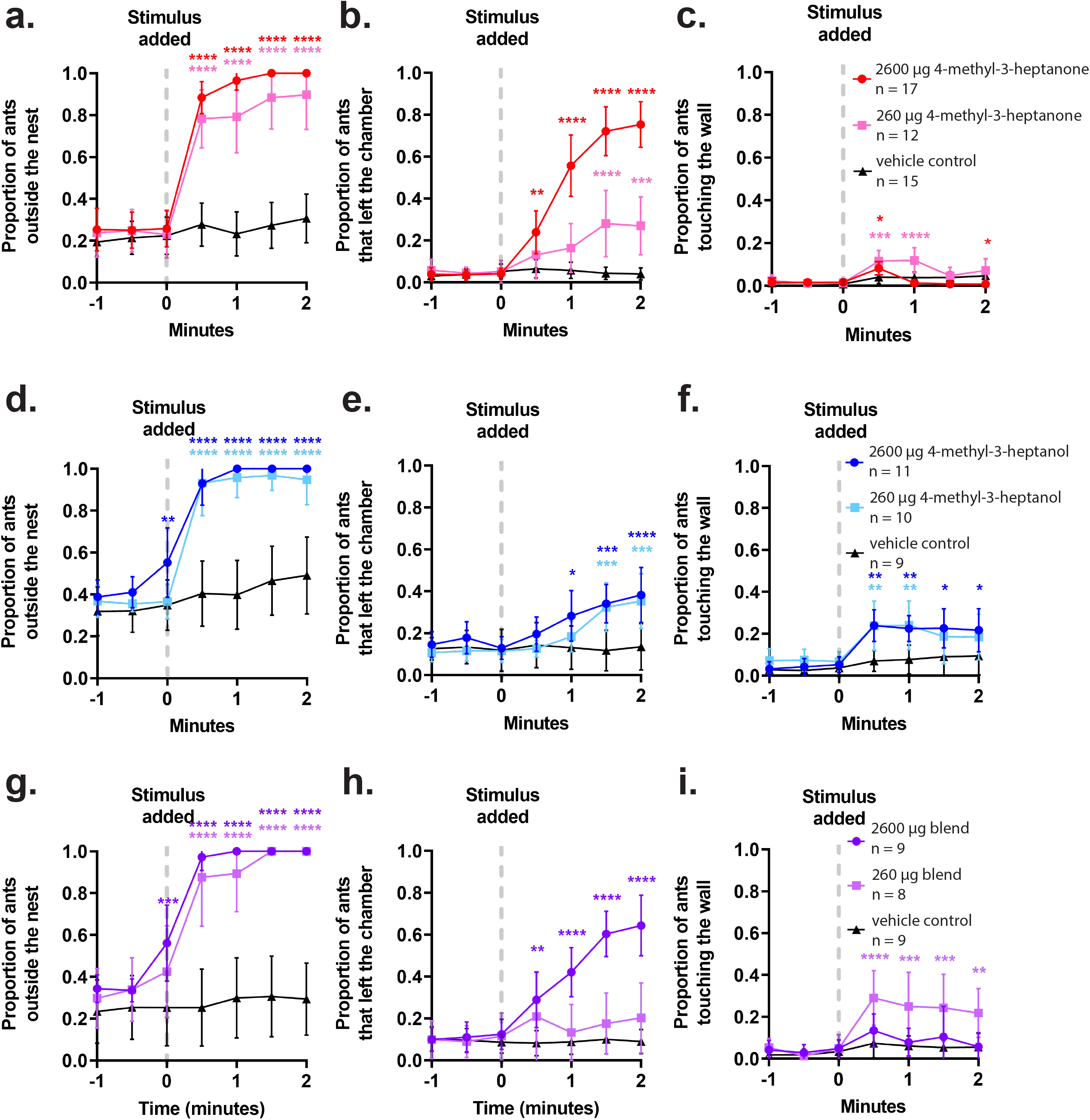
Behavioral response to candidate alarm pheromone components. Quantification of features of the behavioral response of *O. biroi* colonies to 4-methyl-3-heptanone (a-c), 4-methyl-3-heptanol (d-f), and a blend of 90% 4-methyl-3-heptanone and 10% 4-methyl-3-heptanol (g-i). Data are included from 1 minute prior to adding the stimulus until 2 minutes after. Individual datapoints indicate means and error bars denote 95% confidence intervals. Sample sizes represent replicate colonies tested. Statistical comparisons were performed using a 2-way repeated measures ANOVA with Dunnett’s multiple comparisons test to compare individual timepoints to the vehicle control. *p<0.05, **p<0.01, ***p<0.001, ****p<0.0001.

In response to 4-methyl-3-heptanol, ants also rapidly left the nest pile (**Fig 3d, Table S3, Video S4**) and left the nest chamber in a dose-dependent manner, although to a lesser extent than in response to 4-methyl-3-heptanone (**Fig 3e, Table S3**). Like the response to 4-methyl-3-heptanone, ants were initially attracted to 4-methyl-3-heptanol. However, both concentrations attracted a higher proportion of ants than 4-methyl-3-heptanone, and the attraction persisted beyond the first minute after exposure (**Fig 3f, Table S3**). The higher proportion of attracted ants, along with the persistence of the attraction and fewer ants leaving the nest chamber, indicates that this compound could be more attractive to the ants compared to 4-methyl-3-heptanone.

To compare behavioral responses to 4-methyl-3-heptanone and 4-methyl-3-heptanol directly, we quantified the area under the curve for the first two minutes after adding the stimulus. As anticipated, there was no difference in ants outside the nest pile between both compounds (**Fig S6a, Table S4**), but 4-methyl-3-heptanol was significantly more attractive to ants than 4-methyl-3-heptanone at both tested doses (**Fig S6c, Table S4**), and at the high dose, 4-methyl-3-heptanone was significantly more repulsive to ants (**Fig S6b, Table S4**). These results indicate that, while both compounds are sufficient to induce the alarm response, there are slight differences in the behavioral responses they trigger.

We created a synthetic blend of 90% 4-methyl-3-heptanone and 10% 4-methyl-3-heptanol to mimic the ratio of the two compounds in the head of *O. biroi*, where 4-methyl-3-heptanone is the major component and 4-methyl-3-heptanol is the most abundant minor component (**Fig. 2, Table 1**). This blend triggered ants to rapidly leave the nest pile at both concentrations tested (**Fig 3g, Table S3, Video S5**). At the high dose, ants were significantly more likely to leave the nest chamber (**Fig 3h, Table S3**) but were not very attracted to the compound mix (**Fig 3i, Table S3**). At the lower dose, however, ants did not leave the nest chamber, but were attracted to the source of the odor (**Fig 3h&i, Table S3**). These results, in combination with the area under the curve analysis (**Fig S6, Table S4**), indicate that there is no obvious synergistic interaction between 4-methyl-3-heptanone and 4-methyl-3-heptanol in the synthetic alarm pheromone blend. Instead, the high dose of the blend is more repulsive, like 4-methyl-3-heptanone, and the low dose is more attractive, like 4-methyl-3-heptanol. While we did not observe any synergistic interaction between 4-methyl-3-heptanone and 4-methyl-3-heptanol, it is possible that this type of interaction occurs at very low doses, where a single compound alone might not be sufficient to induce a behavioral response.

## DISCUSSION

In this study, we characterized the alarm behavior of the clonal raider ant, *O. biroi*, and identified the chemical components of its alarm pheromone. The alarm response of *O. biroi* is characteristic of a panic alarm response, with ants becoming unsettled, leaving the nest, and moving away from the source of alarm. Alarm pheromone is released from the head of the ant, and we identified two volatile compounds as candidate alarm pheromone components. These two compounds, 4-methyl-3-heptanone and 4-methyl-3-heptanol, are known alarm pheromones in other ant species, are detected by the antennae of *O. biroi*, and are sufficient to trigger a behavioral alarm response, both alone and in combination. These results suggest that the alarm pheromone of *O. biroi* includes a blend of 4-methyl-3-heptanone and 4-methyl-3-heptanol. Future studies identifying the compounds released by alarmed ants will provide additional insight into the exact chemical composition of the alarm pheromone and whether there are minor components found in the head or elsewhere in the body that act to modulate the behavioral response to the major compounds tested here.

In cases where alarm pheromones in ants are released from the head, the mandibular gland is often the source (Wood et al. 2011). 4-methyl-3-heptanone and 4-methyl-3-heptanol have been found together in the mandibular glands or heads of other ants, including some species of *Eciton* army ants that are relatives of *O. biroi* in the ant subfamily Dorylinae (Riley et al. 1974; Pasteels et al. 1981; Hernández et al. 1999; Bento et al. 2007; Brückner et al. 2018). Together, this suggests that 4-methyl-3-heptanone and 4-methyl-3-heptanol are likely released from the mandibular gland in *O. biroi*. However, due to the small size of these ants, we were unable to verify this experimentally by extracting mandibular contents directly.

We studied behavioral responses of clonal raider ant colonies to two different doses of the synthetic alarm pheromone compounds, 260 μg and 2600 μg. While these amounts are substantially larger than the amount of each compound found in a single ant, we do not know the biologically relevant amount of compound the ants were exposed to in the behavioral arena. To prevent the pentane solvent from inducing a behavioral effect, we left the diluted compounds and controls to evaporate for 30 seconds on filter paper before exposing the ants. While 4-methyl-3-heptanone and 4-methyl-3-heptanol are less volatile than pentane, they are still quite volatile and some of the compounds evaporated during that time. Furthermore, stereochemistry is important for biological activity in many pheromones (Mori 2007), and both 4-methyl-3-heptanone and 4-methyl-3-heptanol are chiral, with 4-methyl-3-heptanone having a single chiral center and 4-methyl-3-heptanol having two chiral centers (Riley and Silverstein 1974; Einterz et al. 1977; Zada et al. 2004). We have not yet identified the biologically relevant stereoisomer(s) used by *O. biroi*, and therefore used synthetic compounds that were a mixture of stereoisomers. It is possible that the activity of 4-methyl-3-heptanone and 4-methyl-3-heptanol in *O. biroi* is dependent on its stereochemistry, adding further uncertainty about the behaviorally relevant amount of compound perceived by the ants during behavioral experiments. Future work will be required to quantify the amount of compound that reaches the ants in our bioassay, and to conduct additional behavioral experiments with doses that more closely approximate what ants would perceive under naturalistic conditions.

The two compounds that make up the alarm pheromone in *O. biroi* elicit similar, though slightly different, behavioral responses at the doses tested here. 4-methyl-3-heptanone leads ants to become unsettled and move away from the compound after a quick initial period of attraction, whereas 4-methyl-3-heptanol also induces ants to become unsettled but is more attractive and less repulsive. In combination, these compounds trigger a dose-dependent behavioral response, where at low concentrations ants are initially attracted to the pheromone, but at high concentrations they are repelled and move away from the source of the compound. In other ant species that use multicomponent alarm pheromones, with components that elicit different behavioral effects, alarm behaviors can depend on the total or relative amounts of each component present in the alarm pheromone (Bradshaw et al. 1975, 1979; Fujiwara-Tsujii et al. 2006). We hypothesize that an individual ant may release more alarm pheromone or other ants in the colony may also release alarm pheromone in response to more urgent or dangerous threats, thereby amplifying the signal and triggering a behavioral response that might better protect the colony.

While 4-methyl-3-heptanone and 4-methyl-3-heptanol have been previously described as alarm pheromones in other ant species, this is the first description of the alarm pheromone and alarm behavior in a non-army ant doryline, and the first identified pheromone for *O. biroi*. Because *O. biroi* can be maintained under standardized laboratory conditions and is genetically accessible (Trible et al. 2017), identification of its alarm pheromone will facilitate future work studying the behavioral, genetic, and neuronal underpinnings of the alarm response in ants.

## STATEMENTS AND DECLARATIONS

This work was supported by the National Institute of General Medical Sciences of the NIH under Award R35GM127007 to D.J.C.K. The content is solely the responsibility of the authors and does not necessarily represent the official views of the NIH. Additional support was provided by a Faculty Scholars Award from the Howard Hughes Medical Institute to D.J.C.K. L.E.L. was supported by an NSF Graduate Research Fellowship under award number DGE 194642. This work was supported in part by a grant to The Rockefeller University from the Howard Hughes Medical Institute through the James H. Gilliam Fellowships for Advanced Study program. T.S. acknowledges the support by the DFG State major instrumentation program and the State of Bavaria, Germany. D.J.C.K. is an investigator of the Howard Hughes Medical Institute. This is Clonal Raider Ant Project paper number 24.

### The authors have no relevant financial or non-financial interests to disclose

All authors contributed to study conception and design. Lindsey E. Lopes performed behavioral experiments and data analysis, with guidance from Daniel J. C. Kronauer. Erik T. Frank performed chemical experiments and data analysis, with guidance from Thomas Schmitt. Erik T. Frank and Zsolt Kárpáti performed the electroantennographic recordings and analyzed the data. Lindsey E. Lopes and Daniel J. C. Kronauer wrote a first draft of the manuscript, and all authors contributed to the final manuscript.

We thank Leonora Olivos Cisneros and Stephany Valdés Rodríguez for assistance with ant maintenance, the Rockefeller University Precision Instrumentation Technologies for assistance with arena construction, Vikram Chandra and Asaf Gal for assistance with arena design and behavioral data analysis, and Yuko Ulrich for providing ants for chemical analyses. We also thank Yuko Ulrich and members of the Laboratory of Social Evolution and Behavior for helpful conversations.

The behavioral datasets and raw mass spectra from this study are available in the Zenodo repository with DOI 10.5281/zenodo.7216951.

## SUPPLEMENTARY INFORMATION

**Figure S1.** Alarm arena design. The alarm arena had two areas separated by a tunnel. Each area consists of a small rectangular stimulus chamber and a large square nest chamber, separated by a mesh wall (denoted by a purple dashed line in the figure). These chambers have separate clear plastic acrylic lids, allowing access to the stimulus chamber without disturbing ants in the nest chamber. The brown circle represents the nest pile, where ants and their eggs are tightly clustered prior to starting the experiment. Created with BioRender.com

**Figure S2.** Full time course of characterization of alarm behavior and localization of alarm pheromone in *O. biroi*. Quantification of features of the behavioral response of *O. biroi* colonies to a live alarmed ant (a-c) and crushed body parts of an ant (d-f). Each datapoint indicates the mean and error bars indicate the 95% confidence interval. Sample sizes represent replicate colonies tested.

**Figure S3.** Full time course of behavioral response to candidate alarm pheromone components. Quantification of features of the behavioral response of *O. biroi* colonies to 4-methyl-3-heptanone (a-c), 4-methyl-3-heptanol (d-f), and a blend of 90% 4-methyl-3-heptanone and 10% 4-methyl-3-heptanol (g-i). Each datapoint indicates the mean and error bars indicate the 95% confidence interval. Sample sizes represent replicate colonies tested.

**Figure S4.** Chemical compounds in the ant body. Gas-chromatographic representation of one sample of 5 pooled workers (a), 5 mesosomas (b) and 5 gasters (c). Compounds found in the head are numbered and can be found in Table 2.

**Figure S5.** Antennal detection of candidate alarm pheromone components. Results from EAG recordings in response to 1 μg, 10 μg, and 100 μg of 4-methyl-3-heptanone or 4-methyl-3-heptanol and the solvent control pentane. In total, 9 antennae were tested, except for the 1 μg 4-methyl-3-heptanone condition where 8 antennae were tested. Statistical comparisons were made using a mixed-effects analysis with a Geisser-Greenhouse correction and Dunnett’s multiple comparisons test to compare the response to each compound with the solvent control. *p<0.05, **p<0.01, ***p<0.001, ****p<0.0001.

**Figure S6.** Comparison of behavioral responses to candidate alarm pheromone components and the synthetic alarm pheromone blend. Area under the curve the first 2 minutes after adding the stimulus for ants outside the nest pile (a), ants repelled from the compound(s) (b), and ants attracted to the compound(s) (c). The two compounds and blend were tested in a separate set of experiments and a vehicle control (in grey) was run for each set of experiments. Each datapoint indicates the mean, and error bars represent the 95% confidence intervals. Statistical comparisons were performed using a 2-way ANOVA with Tukey’s multiple comparisons tests to compare the different compounds and blend across concentrations. *p<0.05, **p<0.01, ***p<0.001, ****p<0.0001.

**Video S1.** Representative videos of the behavioral response to control (top) and a live alarmed nestmate (bottom)“. The initial nest pile is in the left nest chamber, and baseline activity was recorded for 5 minutes. 5 minutes into the recording, the stimulus (live alarmed ant or paper control) is added to the stimulus chamber on the right side. The video is sped up 8x, and addition of the stimulus is indicated by a red circle in the top right corner.

**Video S2.** Representative videos of the behavioral response to a crushed body (top) and a crushed head (bottom). The initial nest pile is in the left nest chamber, and baseline activity was recorded for 5 minutes. 5 minutes into the recording, the stimulus (crushed head or crushed body) is added to the stimulus chamber on the right side. The video is sped up 8x, and addition of the stimulus is indicated by a red circle in the top right corner.

**Video S3.** Representative videos of the behavioral response to the vehicle control (top) and two amounts of 4-methyl-3-heptanone, 260 μg (middle) and 2600 μg (bottom). The initial nest pile is in the left nest chamber, and baseline activity was recorded for 5 minutes. 5 minutes into the recording, the stimulus (filter paper with 2600 μg 4-methyl-3-heptanone, 260 μg 4-methyl-3-heptanone, or vehicle control) is added to the stimulus chamber on the right side. The video is sped up 8x, and addition of the stimulus is indicated by a red circle in the top right corner.

**Video S4.** Representative videos of the behavioral response to the vehicle control (top) and two amounts of 4-methyl-3-heptanol, 260 μg (middle) and 2600 μg (bottom). The initial nest pile is in the left nest chamber, and baseline activity was recorded for 5 minutes. 5 minutes into the recording, the stimulus (filter paper with 2600 μg 4-methyl-3-heptanol, 260 μg 4-methyl-3-heptanol, or vehicle control) is added to the stimulus chamber on the right side. The video is sped up 8x, and addition of the stimulus is indicated by a red circle in the top right corner.

**Video S5**. Representative videos of the behavioral response to the vehicle control (top) and two amounts of a blend of 90% 4-methyl-3-heptanone and 10% 4-methyl-3-heptanol, 260 μg (middle) and 2600 μg (bottom). The initial nest pile is in the left nest chamber, and baseline activity was recorded for 5 minutes. 5 minutes into the recording, the stimulus (filter paper with 2600 μg blend, 260 μg blend, or vehicle control) is added to the stimulus chamber on the right side. The video is sped up 8x, and addition of the stimulus is indicated by a red circle in the top right corner.

**Table S1.** Statistical analysis of characterization of alarm behavior and localization of alarm pheromone. Table includes the statistical analyses from the quantification of features of the behavioral response of *O. biroi* colonies to a live alarmed ant and crushed body parts of an ant. Statistical comparisons were performed using a 2-way RM ANOVA with Šidák’s multiple comparisons test to compare individual timepoints.

| Experiment | Behavior | Source of variation<br>(Two-way RM ANOVA) |  |  | Number of<br>arenas |
| --- | --- | --- | --- | --- | --- |
|  |  | Time x<br>Stimulus | Time | Stimulus |  |
| Characterizing<br>alarm behavior<br>(Fig 1a-c) | Outside nest pile | 19.48%<br>p < 0.0001 | 24.35%<br>p < 0.0001 | 12.87%<br>p = 0.0009 | Alarmed ant<br>n = 13 |
|  | Left nest chamber | 13.71%<br>p < 0.0001 | 24.90%<br>p < 0.0001 | 6.196%<br>p = 0.0280 | Control paper<br>n = 10 |
|  | Touching wall | 2.991%<br>p = 0.2342 | 1.872%<br>p = 0.5310 | 3.845%<br>p = 0.1928 |  |
| Localization of<br>alarm pheromone<br>(Fig 1d-f) | Outside nest pile | 11.75%<br>p < 0.0001 | 15.22%<br>p < 0.0001 | 29.19%<br>p = 0.0004 | Crushed head<br>n = 11 |
|  | Left nest chamber | 8.380%<br>p = 0.0001 | 8.013%<br>p = 0.002 | 10.40%<br>p = 0.0329 | Crushed body<br>n = 11 |
|  | Touching wall | 9.666%<br>p < 0.0001 | 13.96%<br>p < 0.0001 | 31.18%<br>p < 0.0001 |  |

**Table S2.** Statistical analysis of EAG recordings. Antennal sensitivity to 1 μg, 10 μg, and 100 μg of 4-methyl-3-heptanone and 4-methyl-3-heptanol and a solvent control were compared using a mixed-effects analysis with a Geisser-Greenhouse correction and Dunnett’s multiple comparisons test was used to compare each compound to the solvent.

| Experiment | Mixed-effects analysis | Compound (compared to solvent) | Adjusted P Value |
| --- | --- | --- | --- |
| Antennal sensitivity to candidate compounds (Fig S5) | Difference between treatments<br>p=0.0022 | 1 μg 4-methyl-3-heptanone | p = 0.9998 |
| | | 10 μg 4-methyl-3-heptanone | p = 0.0451 |
| | | 100 μg 4-methyl-3-heptanone | p = 0.0112 |
| | | 1 μg 4-methyl-3-heptanol | p = 0.9085 |
| | | 10 μg 4-methyl-3-heptanol | p = 0.0713 |
| | | 100 μg 4-methyl-3-heptanol | p = 0.0100 |

**Table S3.** Statistical analysis of behavioral responses to candidate alarm pheromone components. Quantification of features of the behavioral response of *O. biroi* colonies to 4-methyl-3-heptanone, 4-methyl-3-heptanol, and a blend of 90% 4-methyl-3-heptanone and 10% 4-methyl-3-heptanol. Statistical comparisons were performed using a 2-way RM ANOVA with Dunnett’s multiple comparisons test to compare individual timepoints to the vehicle control.

| Experiment | Behavior | Source of variation<br>(Two-way RM ANOVA) |  |  | Number of arenas |
| --- | --- | --- | --- | --- | --- |
|  |  | Time x Stimulus | Time | Stimulus |  |
| Response to 4-methyl-3-heptanone (Fig 3a-c) | Outside nest pile | 15.37%<br>p < 0.0001 | 37.71%<br>p < 0.0001 | 24.87%<br>p < 0.0001 | 2600 µg n = 17 |
|  | Left nest chamber | 23.88%<br>p < 0.0001 | 23.06%<br>p < 0.0001 | 22.13%<br>p < 0.0001 | 260 µg n = 12 |
|  | Touching wall | 10.59%<br>p < 0.0001 | 16.80%<br>p < 0.0001 | 6.696%<br>p = 0.0089 | control n = 15 |
| Response to 4-methyl-3-heptanol (Fig 3d-f) | Outside nest pile | 11.29%<br>p < 0.0001 | 40.97%<br>p < 0.0001 | 24.62%<br>p < 0.0001 | 2600 µg n = 11 |
|  | Left nest chamber | 9.072%<br>p < 0.0001 | 16.99%<br>p < 0.0001 | 8.514%<br>p = 0.0510 | 260 µg n = 10 |
|  | Touching wall | 4.923%<br>p = 0.0537 | 22.44%<br>p < 0.0001 | 9.693%<br>p = 0.0103 | control n = 9 |
| Response to blend (Fig 3g-i) | Outside nest pile | 13.11%<br>p < 0.0001 | 31.13%<br>p < 0.0001 | 34.56%<br>p < 0.0001 | 2600 µg n = 9 |
|  | Left nest chamber | 22.31%<br>p < 0.0001 | 16.44%<br>p < 0.0001 | 24.55%<br>p < 0.0001 | 260 µg n = 8 |
|  | Touching wall | 9.727%<br>p = 0.0006 | 17.46%<br>p < 0.0001 | 14.72%<br>p = 0.0037 | control n = 9 |

**Table S4.** Statistical analysis of area under the curve 2 minutes following exposure to candidate alarm pheromone components and the blend. Comparison of 4-methyl-3-heptanone, 4-methyl-3-heptanol, and 90% 4-methyl-3-heptanone / 10% 4-methyl-3-heptanol blend in ants outside the nest pile, ants repelled from the compound(s), and ants attracted to the compound(s). Statistical comparisons were performed using a 2-way ANOVA with Tukey’s multiple comparisons tests to compare the different compounds and blend across concentrations.

| Behavior | Source of variation<br>(Two-way ANOVA) |  |  |
| --- | --- | --- | --- |
|  | Concentration x<br>Compound | Concentration | Compound |
| Ants outside the nest pile<br>– unsettled<br>(Fig S4a) | 0.6611%<br>p = 0.7941 | 57.52%<br>p < 0.0001 | 2.072%<br>p = 0.0775 |
| Ants that left the nest<br>chamber – repulsion<br>(Fig S4b) | 5.335%<br>p = 0.1084 | 26.49%<br>p < 0.0001 | 0.3398%<br>p = 0.7803 |
| Ants that are touching the<br>wall – attraction<br>(Fig S4c) | 6.088%<br>p = 0.0967 | 11.58%<br>p = 0.0008 | 14.41%<br>p = 0.0002 |

